# Nona: A unifying multimodal masking framework for functional genomics

**DOI:** 10.1101/2025.11.06.687036

**Authors:** Surag Nair, Ehsan Hajiramezanali, Alex Tseng, Nathaniel Diamant, Johannes Hingerl, Avantika Lal, Tommaso Biancalani, Héctor Corrada Bravo, Gabriele Scalia, Gökcen Eraslan

## Abstract

The non-coding genome encodes complex regulatory logic that orchestrates gene expression and cell identity. While machine learning models for functional genomics have advanced our understanding of the cis-regulatory code, sequence-to-function models, DNA language models, and generative models have evolved as separate paradigms despite probing the same underlying regulatory biology. We introduce Nona, a multimodal masked modeling framework that unifies these paradigms by learning jointly from DNA sequence and base-resolution functional genomics data. Beyond unifying existing modeling paradigms, Nona enables entirely new modeling objectives. We demonstrate its versatility through three applications: (1) a context-aware sequence-to-function model that improves local predictions by up to 13% by correcting systematic errors in sequence-to-function predictions; (2) a functional language model that integrates functional data into language modeling, learns relevant regulatory sequence motifs, and enables regulatory element design through masked discrete diffusion; (3) functional genotyping, which reveals an unrecognized privacy vulnerability in processed ATAC-seq data and re-identifies individuals from genetic databases with perfect accuracy. Together, these results establish masking as a universal interface for integrated modeling of functional genomics data, unifying disparate approaches while opening new directions for understanding and engineering the regulatory genome.

## Introduction

The non-coding genome encodes a complex regulatory logic that orchestrates gene expression, cell identity, developmental trajectories, and organismal homeostasis. Understanding this code remains a fundamental challenge in biology. Machine learning models for genomics have greatly advanced the understanding of this code by improving our ability to predict functional genomics signals, the consequences of genetic variation, and to edit or design DNA sequences^1^. Three dominant paradigms have evolved. Sequence-to-function models, which predict functional genomics tracks such as DNase-seq, ChIP-seq, CAGE-seq, and RNA-seq from DNA sequence, learn the sequence logic underlying transcription factor binding, long-range regulatory interactions, splicing, and ultimately, transcriptional regulation^2–4^. These models represent the state of the art for predicting the effects of variants on chromatin accessibility and gene expression^5^. More recently, DNA language models trained purely on sequence data have demonstrated the predictive power contained in co-evolutionary sequence signals^6–9^. In parallel, conditional generative models have expanded our ability to design realistic regulatory DNA sequences with desired activities across cell states^10–12^.

While extremely powerful, these dominant paradigms are rigid. Despite probing the same underlying regulatory biology, supervised sequence-to-function models, self-supervised language models, and generative models have evolved in isolation, each through narrow, specialized interfaces optimized for a single prediction direction. They do not fully leverage the multimodal nature of DNA sequence and functional genomics data, which are inherently anchored to a common genomic coordinate system. For example, DNA language models lack functional grounding: while they learn sequence features over-represented across the genome and over evolution, they are unable to connect them to cell-type specific regulation. Conversely, conditioning sequence-to-function predictions on additional functional tracks remains non-trivial^13–15^. This fragmentation limits our ability to build models that reason flexibly across sequence and function.

Multimodal masked modeling has emerged as a powerful framework for unifying diverse data modalities. Models that learn by predicting masked inputs across paired data, such as image– text pairs^16,17^, or protein sequence-structure-function representations^18^, have demonstrated how a single architecture can support expressive, flexible learning and enable new applications.

Here, we introduce Nona, a multimodal masked model that operates jointly on DNA sequence and base-resolution functional tracks. Nona takes in DNA sequence and tracks, one or both of which can be partially or fully masked, and predicts the masked values. Through a flexible masking scheme, Nona can be configured for a wide range of tasks, unifying sequence-to-function prediction and language modeling while opening up entirely new classes of applications.

We demonstrate three novel applications of Nona. The first extends sequence-to-function modeling by conditioning on experimental measurements from adjacent regions to improve local functional predictions. The second extends DNA language modeling by conditioning on functional tracks, yielding a “functional language model” that learns transcriptionally relevant sequence features and can also serve as a discrete diffusion model that can design sequences with desired activity. The third reveals an unrecognized privacy risk: a small Nona model trained on base-resolution ATAC-seq data shows that ATAC-seq fragment files can leak private genetic information despite lacking explicit variant information.

These applications illustrate how a unified framework can address disparate challenges in regulatory genomics, while revealing unexpected vulnerabilities in standard data formats. Together, Nona establishes masking as a universal interface for probing, interpreting, and designing the non-coding genome.

### Nona Overview

Nona is built on a simple principle: any genomic modeling task can be formulated as predicting masked values from unmasked context. This unifying view allows diverse applications to be expressed through masking configurations rather than specialized architectures.

At its core, Nona takes in masked DNA sequence and tracks, together with binary matrices indicating which positions and tracks are masked, and predicts the masked values (Fig. 1a,b, Supp Fig. 1a, Methods). Masked positions are set to 0 in the input, and the binary masks allow the model to distinguish true zeros from masked zeros. The model jointly processes all modalities to predict masked values, learning a shared representation that captures the relationship between sequence and function (Methods).

**Figure 1:**
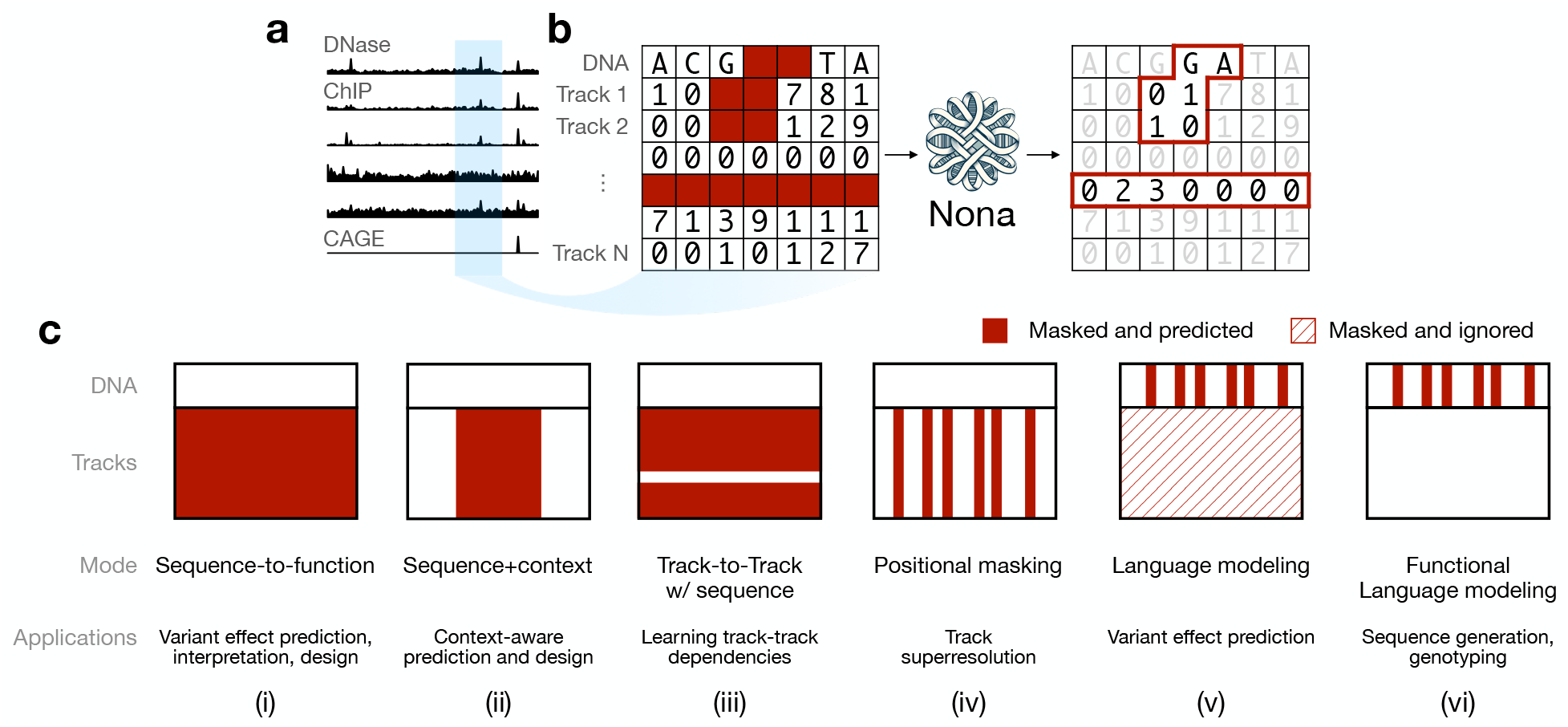
The Nona multimodal masked modeling framework. **a)** Example functional genomics tracks used as base-resolution inputs alongside DNA sequence **b)** Conceptual schematic of Nona’s multimodal masked modeling framework. DNA sequence and multiple functional tracks are provided as input, with selected positions masked according to a specified masking scheme. **c)** Representative masking configurations and their corresponding applications: (i) sequence-to-function, (ii) context-aware prediction, (iii) track-to-track, (iv) positional masking, (v) language modeling, and (vi) functional language modeling. Masked values (red) are predicted, while masked-and-ignored positions (hatched) are excluded from loss computation. By varying the masking configuration, Nona can perform a wide range of tasks spanning prediction, inference, and generation. This work demonstrates applications i, ii, v and vi.

During training, a sequence and corresponding tracks are retrieved, and an input mask is sampled from a pre-specified distribution (Fig. 1c). The input sequence and tracks are multiplied with this mask to zero out masked positions. A separate output mask specifies which positions the loss is computed on. Often, the output mask is the inverse of the input mask (Supp Fig. 1b). The power of this approach lies in its flexibility. This formulation encompasses many diverse configurations, including:

- Sequence-to-function: sequence unmasked; tracks masked (Fig. 1c(i))
- Context-aware: tracks masked in a central focal interval, unmasked elsewhere; sequence unmasked (Fig. 1c(ii))
- Track-to-track: a subset of tracks and sequence unmasked; remaining tracks masked (Fig. 1c(iii))
- Positional masking: a subset of positions masked across all tracks; rest unmasked (Fig. 1c(iv))
- Language modeling: sequence masked at a subset of positions; tracks masked and not predicted (Fig. 1c(v))
- Functional language modeling: sequence masked at a subset of positions; tracks unmasked (Fig. 1c(vi))

For tracks, we use a Poisson negative log-likelihood loss. For sequence, we use a cross entropy loss, averaged across positions specified by the output mask. A hyperparameter controls the relative weights of the two losses. Nona uses a hybrid convolution-transformer U-Net architecture inspired by Borzoi^2^, with separate convolutional encoders for sequence and tracks (Methods). This design efficiently handles the different statistical properties of discrete sequence and continuous track data while enabling cross-modal learning. The exact parameterization of the architecture differs by application.

The versatile masking approach opens up entirely new classes of applications beyond sequence-to-function prediction and language modeling. We next demonstrate three such applications that showcase Nona’s ability to address diverse challenges in regulatory genomics.

### Context-aware epigenetic predictions

Sequence-to-function models discard experimental track data during inference. We hypothesize that explicitly providing track information as input during both training and inference can improve local functional predictions. We introduce a Nona model that extends sequence-to-function modeling by incorporating measured experimental tracks flanking a focal region. We initialize this “context-aware” model by first training a standard sequence-to-function model (Fig. 2a). This model is then fine-tuned to predict tracks in a smaller focal region from DNA sequence together with measured tracks in surrounding regions, similar to inpainting applications in computer vision^19^.

**Figure 2:**
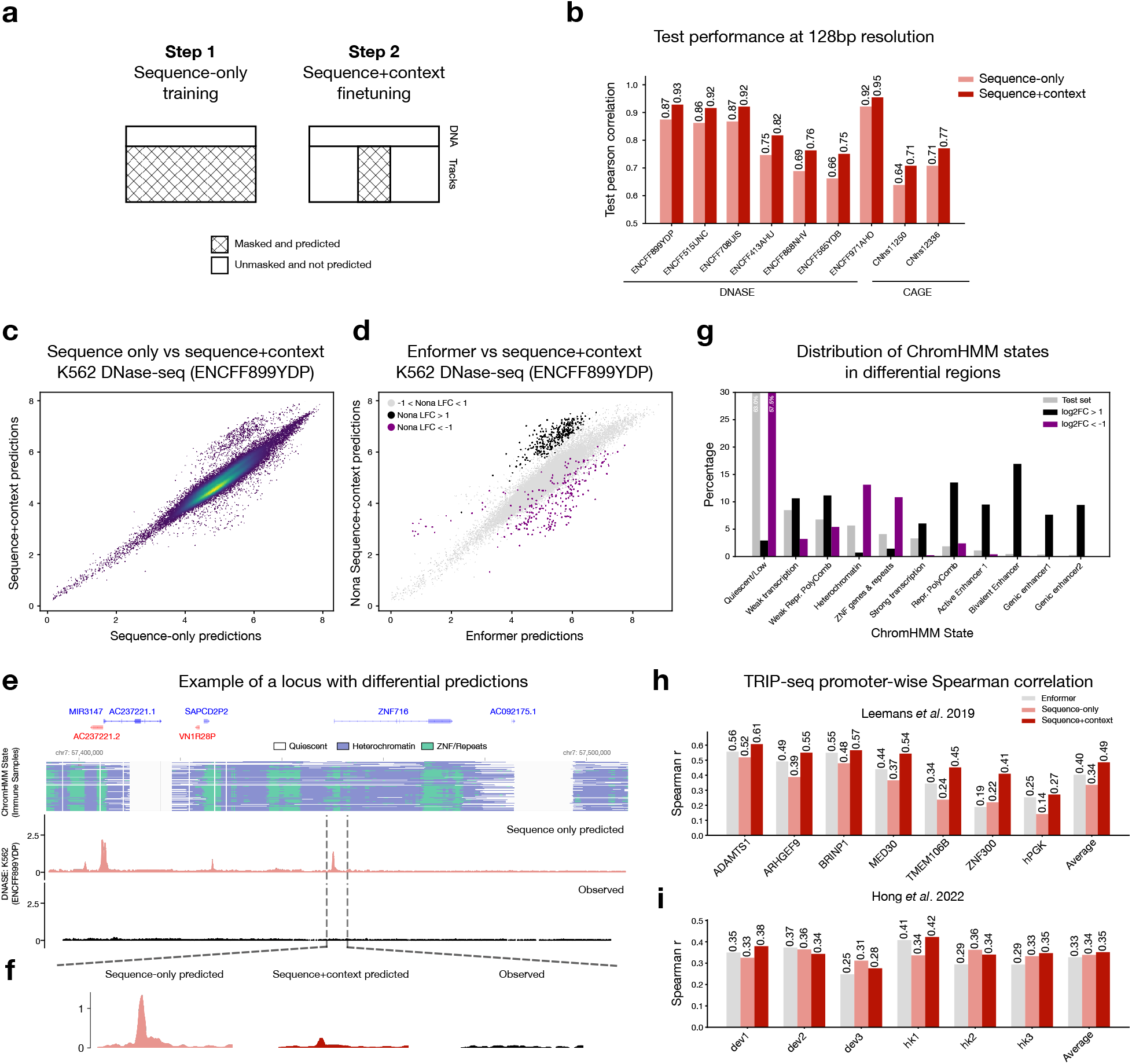
Context-aware sequence modeling improves local predictions by correcting systematic errors in sequence-to-function predictions. **a)** Schematic of the two-stage training procedure: initial sequence-to-function training followed by context-aware fine-tuning using flanking track measurements **b)** Pearson correlation between log1p-transformed predictions and observed values for each human track at 128 bp resolution. Bars show mean across five model replicates. **c)** Comparison of log1p-transformed predictions from the sequence-only and sequence+context models for a representative K562 DNase-seq track. Each point corresponds to a 4096 bp region. Predictions are ensembled across the five replicates. **d)** Same as c) but comparing Enformer predictions with Nona sequence+context predictions. Highlighted points indicate regions with log-fold change (LFC) >1 or < -1 in c). **e)** Example test locus (chr7:57,393,464–57,508,152) showing sequence-only model predictions and observed data. ChromHMM annotations for immune samples are visualized using Epilogos^23^. **f)** Zoomed view of central 4096 bp showing context-aware model correctly predicts low signal by incorporating flanking information **g)** Distribution of ChromHMM states among regions with differential predicted accessibility (log2-fold change >1 or <-1) when comparing sequence+context predictions against both sequence-only and Enformer predictions **h)** Promoter-wise Spearman correlation of model predictions for the Leemans *et al*. TRIP-seq dataset. Correlations are averaged across five replicates for the Nona models. **i)** Same as h) for the Hong *et al*. dataset.

To demonstrate this mode, we trained a model focused on K562 tracks: 7 DNase-seq and 2 CAGE-seq tracks. The model takes a 196608 bp sequence input. In the initial sequence-to-function training, the model predicts the tracks for the central 114688 bp region at base-resolution. In the context-aware fine-tuning, the model predicts tracks in central 4096 bp using the same 196608 bp input sequence along with measured tracks in 96256 bp on either side of the central region. In addition to the human tracks, we also added 4 DNase-seq and 3 CAGE-seq tracks from mouse (Methods). The flexible masking capabilities of Nona seamlessly enable multi-species training (Supp Fig. 2a, Methods). We trained 5 model replicates initialized with different random seeds, using the same training split as the Enformer model^20^.

We evaluated both the sequence-only and fine-tuned context-aware models on the test set. We partitioned the test regions into non-overlapping 4096 bp windows, and extracted the predictions corresponding to these windows for a fair comparison (Methods). Metrics and downstream analyses were performed after aggregating predictions to 128 bp resolution. Because the context-aware model has access to strictly more information than the sequence-only model, we expected its performance to be at least as good. Indeed, the context-aware model achieved higher test set Pearson correlations across all human tracks, with improvements of up to 13% (Fig. 2b).

To investigate what drives this improvement, we compared predicted total counts for one of the DNase-seq tracks across 4096 bp bins (Fig. 2c). While most predictions were highly correlated between the two models, we observed distinct outliers where the context-aware model predicted markedly higher or lower counts. To determine whether these regions correspond to systematic errors in conventional sequence-to-function models, we obtained predictions from Enformer (Methods). Enformer uses the same input and output lengths as the Nona sequence-only model but is trained on a larger set of tracks. We found that these outlier regions were also systematically over- or under-predicted by Enformer relative to the context-aware model (Fig. 2d, Supp Fig. 2b), a pattern that extended to CAGE-seq tracks as well (Supp Fig. 2c, d, e).

At an exemplar locus on chr7, the sequence-only model predicts high DNase-seq activity across the focal and surrounding regions, despite the locus being marked by heterochromatin with no visible DNase-seq peaks (Fig. 2e,f). In contrast, the context-aware model correctly predicts low activity in the focal region by incorporating low signal in the flanking regions. Analysis of chromatin states across all test regions with differential predictions between the two models revealed that these outlier loci are associated with specific chromatin states (Fig. 2g). Regions predicted to have higher activity by the context-aware model are mostly associated with *Repressed Polycomb* and *Enhancer* states, whereas those predicted to have lower activity are linked to *Heterochromatin* and *ZNF genes/repeats* states. These results indicate that sequence-only models systematically mis-predict activity at loci with atypical chromatin states, and that conditioning on local context can correct these errors.

To assess whether context-aware predictions improve performance on downstream tasks, we evaluated the models on two TRIP-seq datasets (Fig. 2h,i). TRIP-seq experiments measure promoter-driven reporter expression across many genomic integration sites, providing an opportunity to test whether incorporating genomic context enhances prediction of reporter activity. On the Leemans *et al*. dataset^21^, the context-aware model consistently outperformed both the sequence-only model and Enformer across different promoters. On the Hong *et al*. dataset^22^, the results were mixed, with the context-aware model performing marginally better on average. Ensembling predictions from model replicates recapitulated these trends (Supp Fig. 2 f,g). Together, these results indicate that context-aware models can improve prediction of local genomic activity.

### Functional language modeling

DNA language models learn powerful representations by predicting masked bases from sequence context alone^6,24^. However, they lack functional grounding: despite learning over-represented patterns and co-evolutionary sequence signals, they cannot connect these features to cell-type specific regulation-a limitation that can be overcome by Nona’s flexible architecture. In this section, we introduce a Nona model to perform a variation of masked language modeling termed “functional language modeling”. This mode involves making predictions from one or many functional tracks in addition to other unmasked bases (Fig. 3a). Functional language modeling corresponds to an input mask where functional tracks are always unmasked, while a fraction of bases are masked, and an output mask that encourages prediction of the masked bases.

**Figure 3:**
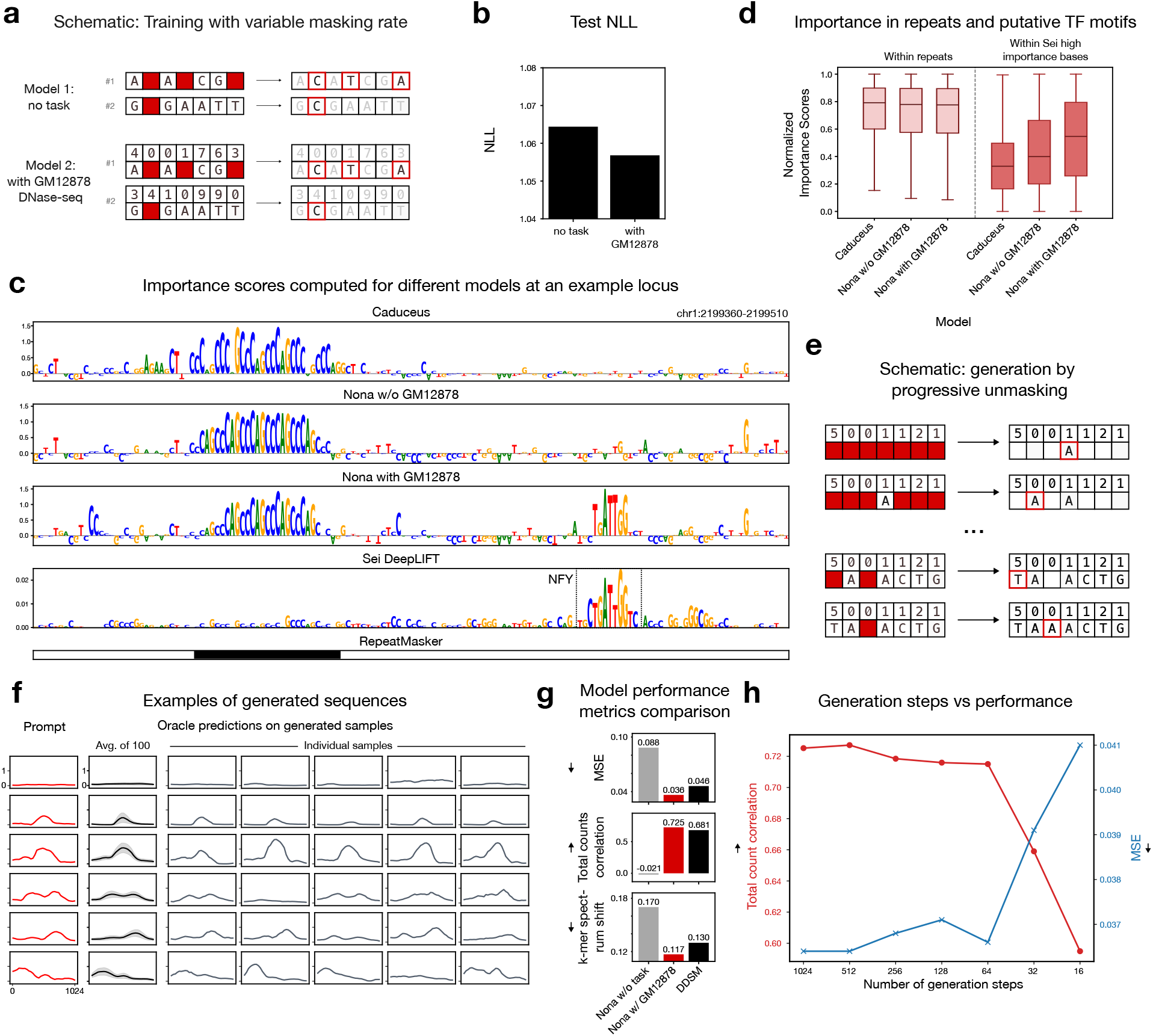
Functional language modeling for interpretation and sequence design. **a)** Schematic of model training with variable masking rate, comparing an unconditional model and a functional language model conditioned on smoothed GM12878 DNase-seq profiles. For both, the rows show two different hypothetical training examples. **b)** Test set negative log-likelihood (NLL) showing improved sequence prediction with functional conditioning (n=46437 sequences, *p*-val<2.2×10^-16^, paired t-test). **c)** Base-level importance scores for different models at an example locus (chr1:2199360-2199510). The conditional Nona model emphasizes a NF-Y transcription factor motif that is not picked up by Caduceus or the unconditional Nona model. **d)** Systematic comparison of normalized importance scores across repeats and putative TF sites identified by Sei. All comparisons paired t-test *p*-val < 2.2 × 10^-16^. **e)** Schematic of sequence generation via progressive unmasking. **f)** Sequence generation examples showing DNase-seq prompts (red, left), averaged oracle predictions over 100 generated sequences (black, middle), and 5 individual sequence predictions (grey, right), demonstrating functional fidelity. The shaded region in the average prediction column indicates the standard deviation across samples. **g)** Generation quality metrics comparing functional Nona (red), unconditional Nona (gray) and DDSM (black). Metrics evaluate functional fidelity (MSE, total count correlation) and sequence realism (k-mer spectrum shift). **h)** Effect of number of generation steps (x-axis, log_2_ scale) on generation quality in terms of total count correlation (y-axis, left) and mean-squared error (MSE, y-axis, right).

We trained a 1024 bp functional language model using GM12878 smoothed DNase-seq as the functional track (Fig. 3a, Methods). Training was performed on DNase-seq peaks and a GC-matched set of non-peak regions. During training, the track was always unmasked, while sequence positions were masked independently with a masking rate sampled from a Uniform(0,1) distribution per example. The model then predicts the masked bases. We also trained a model without functional tracks, serving as an unconditional baseline (Fig. 3a).

The functional model achieved a lower test negative log-likelihood (Fig. 3b, paired t-test *p*-val<2.2×10^-16^, Supp Fig. 3a) and marginally higher base prediction accuracy (Supp Fig. 3b) compared to the unconditional model. To study what additional sequence features were learned by the conditional model, we computed model importance scores (Methods). We initially masked one base at a time and computed the negative entropy of predictions^25^, but found that single-pass inference without any masking produced nearly identical scores, which we used going forward (Supp Fig. 3c). In addition to the Nona models, we also computed importance scores using Caduceus^26^, a masked language model trained on the entire human genome (Methods). To contrast these scores with those obtained from a sequence-to-function model, we generated DeepLIFT/DeepSHAP scores^27^ on the GM12878 DNase-seq outputs of the Sei model^28^ (Methods).

At an exemplar locus on chr1, all language models showed high importance over a simple repeat identified by RepeatMasker^29^ (Fig. 3c). In contrast, Sei showed high importance primarily over an NF-Y motif. Intriguingly, the conditional Nona model, but not the unconditional Nona model, highlighted the same NF-Y motif. This suggests that conditioning on functional data emphasizes transcription factor (TF) motif features.

To test this systematically, we computed importance scores over GM12878 peak sequences in the Nona test set (Methods). Bases overlapping RepeatMasker were labeled as repeats; bases in the top 1% of Sei scores and not in repeats were labeled as putative TF sites. Within repeat regions, all three language models had similar importance (median 0.78) (Fig. 3d). Within TF sites, however, the conditional Nona scored higher (median 0.54) than the unconditional Nona (0.40), which in turn scored higher than Caduceus (0.33, all comparisons paired t-test *p*-val < 2.2 × 10^-16^).

These results demonstrate that in the absence of functional grounding, DNA language models primarily learn easy to identify repeats within non-coding DNA, largely missing TF binding sites. Functional language models capture regulatory features much more strongly than unconditional models. The higher scores of unconditional Nona relative to Caduceus suggest that simply training on cell-type specific regulatory regions already promotes learning of TF features.

### Regulatory element design: a masked discrete diffusion approach

Having shown that functional language models learn regulatory sequence features, we next asked whether they could be used not only for interpretation but also for design. This task can be viewed as a special case of discrete diffusion modeling, where masking acts as the corruption strategy and iterative denoising corresponds to filling in masked positions^30,31^. We implemented a progressive unmasking design strategy in which the model samples bases step by step until the entire sequence is filled (Fig. 3e, Methods). At the start, all positions are masked. At each round, the model predicts base probabilities at every masked position. These probabilities are divided by a temperature parameter T, which sharpens or flattens the distribution; we typically used T=0.7. A random subset of positions is then selected and filled by sampling from the temperature-scaled distribution. In the simplest case, we begin by filling one base at a time. Filled positions remain fixed for the remainder of the process. Repeating this cycle until all positions are filled yields a designed sequence.

As an example, we conditioned on held-out GM12878 DNase-seq profiles and generated sequences *de novo*. Visual inspection of oracle predictions using Borzoi^2^ on multiple generated sequences showed close agreement with the input prompts, recapitulating the shape and scale of the specified profiles (Fig. 3f, Methods). To quantify performance, we compared generated sequences to their matched genomic sequences using a suite of metrics that assess both functional fidelity and sequence realism^11^ (Methods). The functional language model achieved lower mean squared error (MSE), higher correlation of total counts, and smaller sequence composition shift compared to DDSM^12^, a discrete diffusion model based on Dirichlet score matching (Fig. 3g). As a control, we generated sequences using the unconditional Nona model, which showed much higher MSE and no correlation of total counts. More extensive comparisons demonstrate that Nona is highly performant at conditional sequence generation (Supp Fig. 3d).

Compared to autoregressive models, a distinct advantage of discrete diffusion models is that they can decode multiple tokens in parallel at each step. To leverage this ability, we examined how faster decoding affects sequence generation quality. Notably, performance was robust even when the number of generation steps was reduced from 1024 to 64 (i.e. 16× faster), with total count correlation remaining high and MSE low (Fig. 3h). These results highlight the power of a diffusion-style approach for regulatory element design, enabling accurate and efficient generation of sequences that satisfy specified functional constraints. Together, these findings establish functional language modeling as a general framework for both interpreting and designing regulatory elements.

### Functional genotyping

Fragment files from ATAC-seq have become a standard format for sharing data, particularly in single-cell applications^32^. These files contain genomic coordinates of aligned fragments and, unlike raw FASTQ or BAM files, are generally assumed to protect privacy by omitting sequence information. Fragment files can be readily converted into base-resolution BigWig tracks that count Tn5 insertion events, which are often used for training machine learning models^3,5^. Base-resolution ATAC-seq profiles are dominated by Tn5’s local sequence preference^3^. We hypothesized that common genetic variation can modulate ATAC-seq profiles without changing overall accessibility. At a locus on chr1, we observed that the ATAC-seq profiles from individuals in the African Functional Genomics Resource (AFGR)^33^ show systematic differences in profile shape depending on genotype (Fig. 4a, Supp Fig. 4a). This observation raises the possibility that an individual’s genotype can be inferred directly from their ATAC-seq profile.

**Figure 4:**
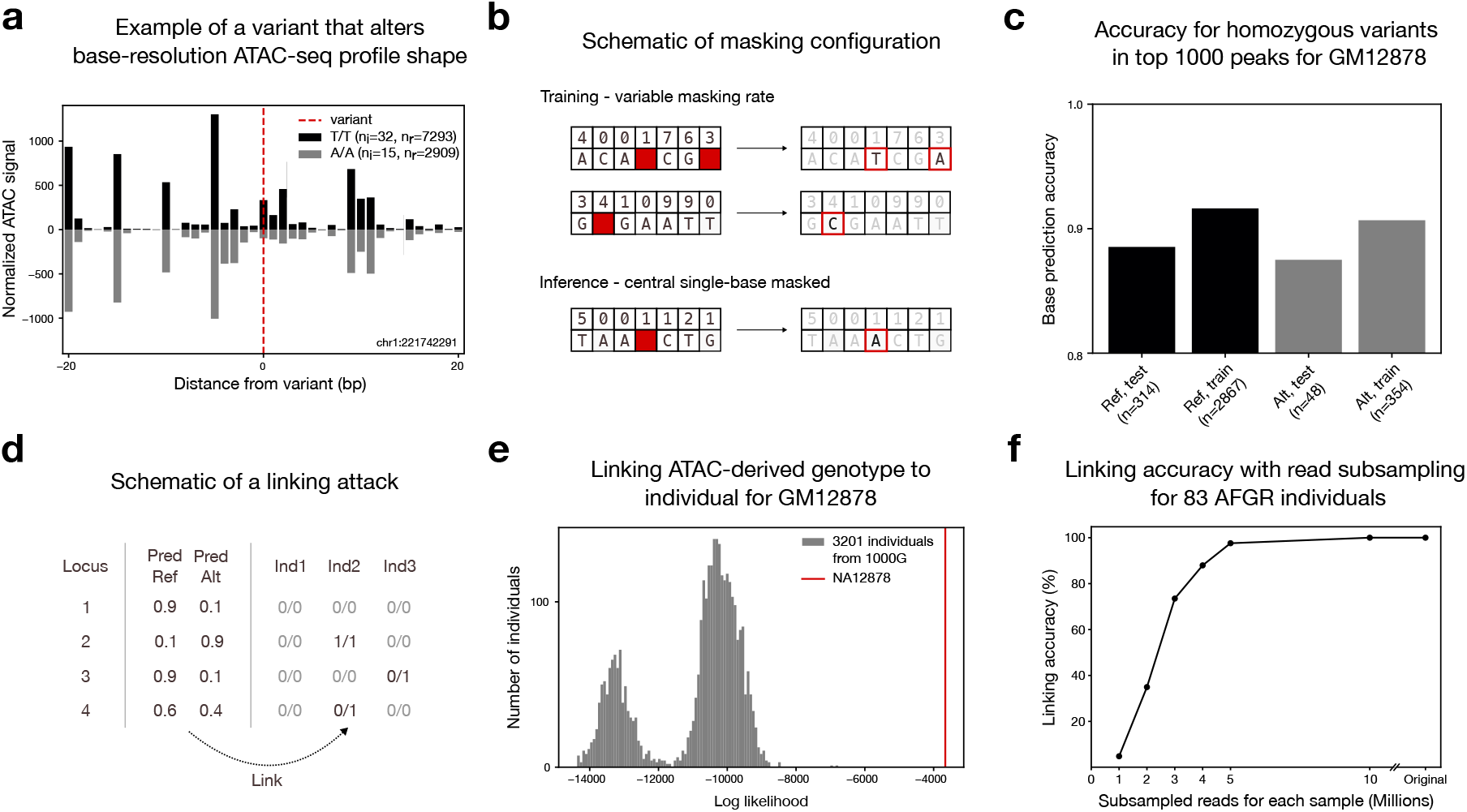
A functional language model trained on base-resolution ATAC-seq profiles enables accurate genotype inference and individual re-identification. **a)** Example of a variant that alters local ATAC-seq profile shape. Signal is shown for individuals from AFGR homozygous for the reference (T/T) or alternate (A/A) allele. The signal is normalized for read depth. The signal for (A/A) allele is reflected along the x-axis. n_i_=number of individuals, n_r_=total number of reads in the window. **b)** Schematic of model training and inference. **c)** Accuracy of base prediction for homozygous variants in GM12878, grouped by allele type and dataset partition. For example, “Ref, train” represents sites where the individual carries the reference allele and the locus was included during training. **d)** Schematic of a linking attack setup. **e)** Distribution of log-likelihood scores from a linking analysis comparing one ATAC-seq sample to 3202 candidate individuals. The red line marks the log-likelihood for the true source individual. **f)** Linking accuracy for 83 individuals from the AFGR dataset as a function of sequencing depth following read subsampling.

To test this, we trained a small 128 bp functional language model on base-resolution GM12878 ATAC-seq (Fig. 4b). As with the DNase-seq functional language model, the track was always unmasked, while sequence positions were masked independently with a masking rate sampled from a Uniform(0,1) distribution per example (Methods). Training used the reference hg38 genome only, with no personal variant information. To evaluate whether the model could predict individual genotype, we obtained NA12878’s (the individual corresponding to GM12878 sample) genotype data for 4756 common SNPs located within the top 1000 strongest LCL ATAC-seq peaks (Methods). At each SNP, we centered the sequence on the variant, masked the focal base, and left the rest of the sequence and track unmasked.

Because the model outputs a single most probable base per position, it cannot easily distinguish heterozygous from homozygous genotypes. We therefore restricted evaluation to sites where NA12878 is homozygous for either the reference or the alternate allele. The model correctly identified the true allele in over 85% of cases (Fig. 4c). Strikingly, performance was equally high at homozygous alternate sites within the training set, even though the model had only been exposed to the reference allele during training and was therefore biased toward predicting the wrong base.

Beyond single-site genotype inference, we next asked whether ATAC-seq profiles could be used to re-identify individuals through a linking attack^34^ (Fig. 4d). In this setting, the input consists of a base-resolution ATAC-seq track from an unknown individual and a panel of candidate genotypes from multiple individuals, one of whom is the true source of the ATAC-seq file. We score each candidate by computing the log-likelihood of their genotypes under the model’s predicted base probabilities at common SNPs (Methods). The individual with the highest score is selected as the predicted source. As a proof of concept, we first applied this method to the GM12878 ATAC-seq data, evaluating it against all 3202 individuals in the 1000 Genomes high-coverage dataset. Using 4756 SNPs within highly accessible regions, NA12878 was unambiguously identified as the top-scoring individual (Fig. 4e).

We next extended this analysis to 83 individuals from AFGR. For each AFGR ATAC-seq sample, we computed linking scores against all 3202 individuals from the 1000 Genomes dataset. Strictly speaking, a separate model should be trained for each sample, as samples differ in terms of quality and sequencing depth. However, we observed that reusing the model trained on GM12878 without any read depth normalization achieved a linking accuracy of 100%, with every sample correctly matched to its source individual. This indicates that the model relies primarily on base-resolution profile shape rather than read depth for genotype inference. Finally, we evaluated robustness to reduced sequencing depth by subsampling the AFGR ATAC-seq data. Linking performance remained near 100% down to 5 million reads per sample, but degraded below this threshold (Fig. 4f). These results demonstrate that base-resolution ATAC-seq profiles are strongly predictive of genotype at common variants, and enable near perfect re-identification of individuals even under modest sequencing coverage.

## Discussion

Nona represents a unified multimodal masked modeling framework for genomic sequence and functional data. By treating masking as a universal interface, Nona connects predictive, generative, and self-supervised learning within a single framework. Rather than developing specialized models for each paradigm, diverse objectives can be expressed through different masking configurations, simplifying how we think about and build models for regulatory genomics. Beyond consolidating existing paradigms, this framing provides a foundation for models that learn a joint latent space of DNA and function, thus enabling reasoning and generation in any direction between the two.

Through its flexible masking scheme, Nona can be configured for a wide range of novel applications. Context-aware modeling revealed that conventional sequence-to-function models systematically under- or over-predict activity in outlier chromatin states, and that such errors can be rescued by conditioning on nearby measured signals. Validation on TRIP-seq data further suggests added value of this masking configuration for predicting expression of integrated reporters and identifying safe harbor or integration sites for cell and gene therapy. However, a limitation of this approach is that context-aware models assume the inserted construct does not disrupt activity in nearby regions.

Functional language modeling bridges the gap between DNA language models and sequence-to-function predictors. While conventional language models excel at capturing co-evolutionary sequence statistics, they are difficult to steer toward functional constraints and often underperform well-tuned sequence-to-function models on downstream tasks^35–37^. Conditioning masked base prediction on functional data provides a route to combine the strengths of both paradigms. Our proof-of-concept model emphasizes transcription factor binding motifs more strongly than unconditional language models. Under variable masking rate regimes, the model is equivalent to a discrete diffusion formulation, which enables efficient, parallel decoding of designed regulatory sequences. More creative masking schemes could guide generation toward more pointed design objectives.

Finally, by training a small functional language model, we demonstrate that base-resolution ATAC-seq signals reveal private genetic information. While the sequence bias of the Tn5 enzyme has been well documented, the ability to invert signals to predict genotype has not been explored^3^. Similar linking attacks have been demonstrated previously in the context of RNA-seq data^34,38^. However, such methods often require a pre-computed panel of eQTLs, or a small cohort of individuals with matched expression and genotypes. In contrast, our method works with a single ATAC-seq sample and just the reference genome. Given the rise in adoption of ATAC-seq, our work recognizes a pertinent and underappreciated privacy risk. These findings have immediate implications for data sharing policies, IRB considerations and public single-cell ATAC-seq repositories. The genomics community should reconsider privacy protections for functional genomics data as linking attacks grow more sophisticated. This may require developing privacy-preserving processing methods that obscure genotype information while preserving biologically relevant patterns such as transcription factor footprints.

Conceptually, Nona provides a straightforward yet powerful consolidation of existing functional genomics model paradigms. Folding different model types into a single framework enhances component reuse across tasks and scales– from the 196 kb context-aware model to the compact 128 bp genotyping model. Looking forward, larger Nona models jointly trained across multiple masking configurations could learn richer shared representations of sequence and function, serving as the next generation of foundation models for regulatory genomics. Together, these results establish Nona as a general-purpose framework for learning and reasoning across sequence and function, unifying prediction and generation under a single modeling principle and opening new directions for the study of the non-coding genome.

## Data availability

All datasets used are publicly available from ENCODE, the 1000 Genomes Project, AFGR, and published ChromHMM/Epilogos annotations. All accession numbers and processing details for every dataset are listed in the Methods section.

## Supplementary Figures

**Supplementary Figure 1:**
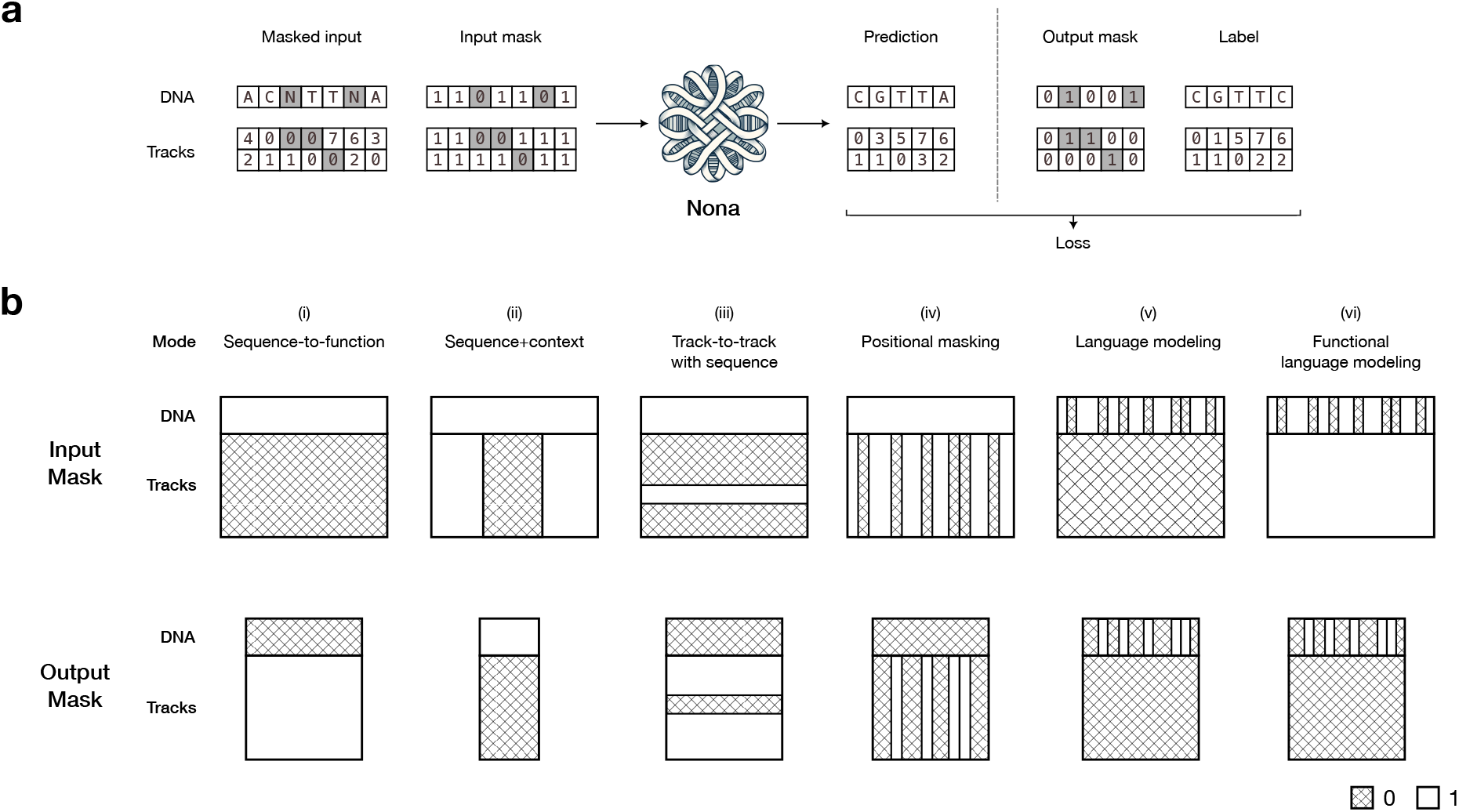
**a)** Conceptual schematic of Nona’s multimodal masked modeling framework in the general case where output length < input length. Masked DNA sequence and multiple functional tracks are provided as input, together with binary matrices defining the input mask. The input mask enables the model to distinguish between true zeros and masked zeros. Nona predicts values at all positions, and the output mask specifies where loss is computed. Shaded squares in the input indicate masked positions; shaded squares in the output mask indicate positions contributing to the loss. **b)** Representative masking configurations from Figure 1c, illustrating a more general case where output length < input length. In the input mask, 1 denotes unmasked values and 0 denotes masked values. In the output mask, 1 marks positions where the loss is computed, and 0 marks positions ignored in the loss. In most cases, the output mask is the inverse of the input mask, although exceptions occur (e.g., mode (v)).

**Supplementary Figure 2:**
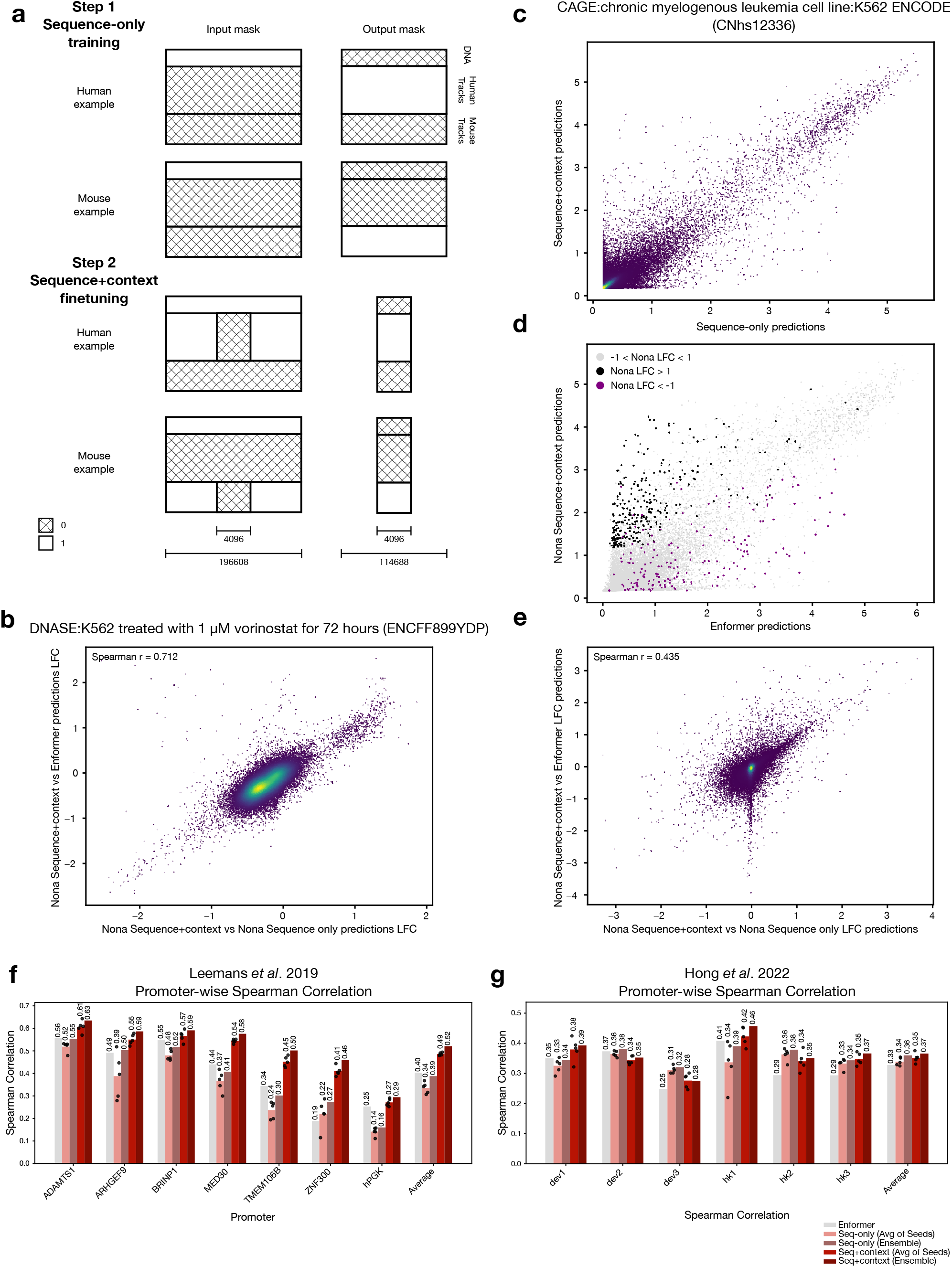
**a)** Schematic of the full two-stage training procedure with human and mouse examples. In the input mask, 1 denotes unmasked values and 0 denotes masked values. In the output mask, 1 marks positions where the loss is computed, and 0 marks positions ignored in the loss. **b)** Comparison of log-fold change (LFC) between Nona sequence+context vs Nona sequence-only predictions and LFC between Nona sequence+context vs Enformer predictions for the DNase-seq track in Fig. 2. **c)** Comparison of log1p-transformed predictions from the sequence-only and sequence+context models for a CAGE-seq track. Each point corresponds to a 4096 bp region. Predictions are ensembled across the five replicates. **d)** Same as c) but comparing Enformer predictions with Nona sequence+context predictions. Highlighted points indicate regions with LFC >1 or < -1 in c). **e)** Same as b) for the CAGE-seq tracks. **f)** Promoter-wise Spearman correlation of model predictions for the Leemans *et al*. TRIP-seq dataset. Correlations are shown for each of the five replicates for the Nona models, as well as for the ensembled predictions. **g)** Same as f) for Hong *et al*. TRIP-seq dataset.

**Supplementary Figure 3:**
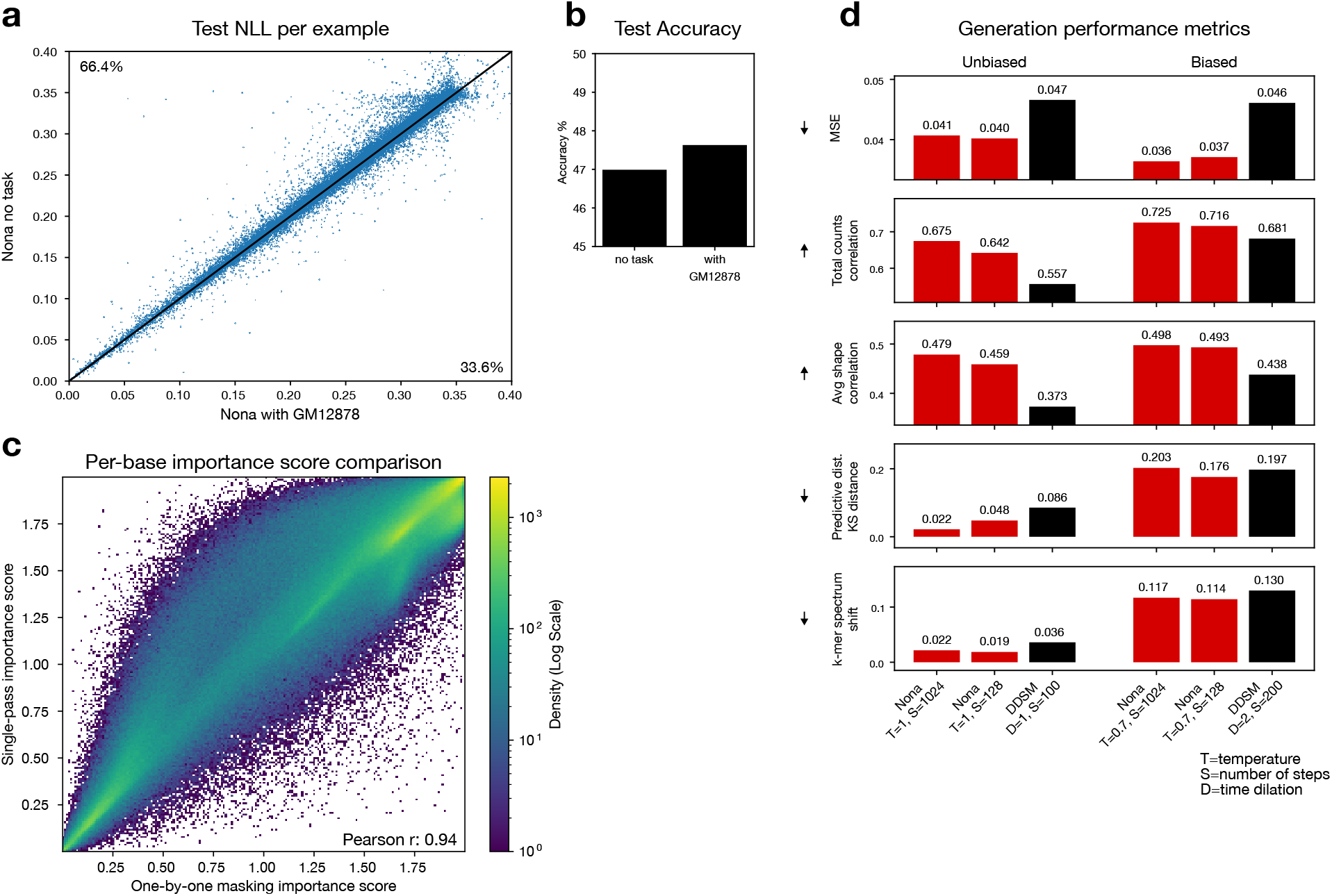
**a)** Related to Fig. 3b, shows test NLL per example comparing the unconditional and functional Nona language models trained on GM12878 DNase-seq. The functional language model has lower test NLL than the unconditional model (n=46437 sequences, *p*-val<2.2×10^-16^, paired t-test) **b)** Test accuracy in terms of bases predicted correctly by the models. **c)** Comparison of per-base importance scores using a one-by-one masking strategy and a single-pass strategy. The one-by-one masking strategy requires as many forward passes through the model as the length of the sequence, while single-pass strategy requires just one. **d)** Extended version of Fig. 3g, showing generation metrics on sequences generated by DDSM (black), and GM12878 DNase-seq functional language model (red) for different generation parameters.

**Supplementary Figure 4:**
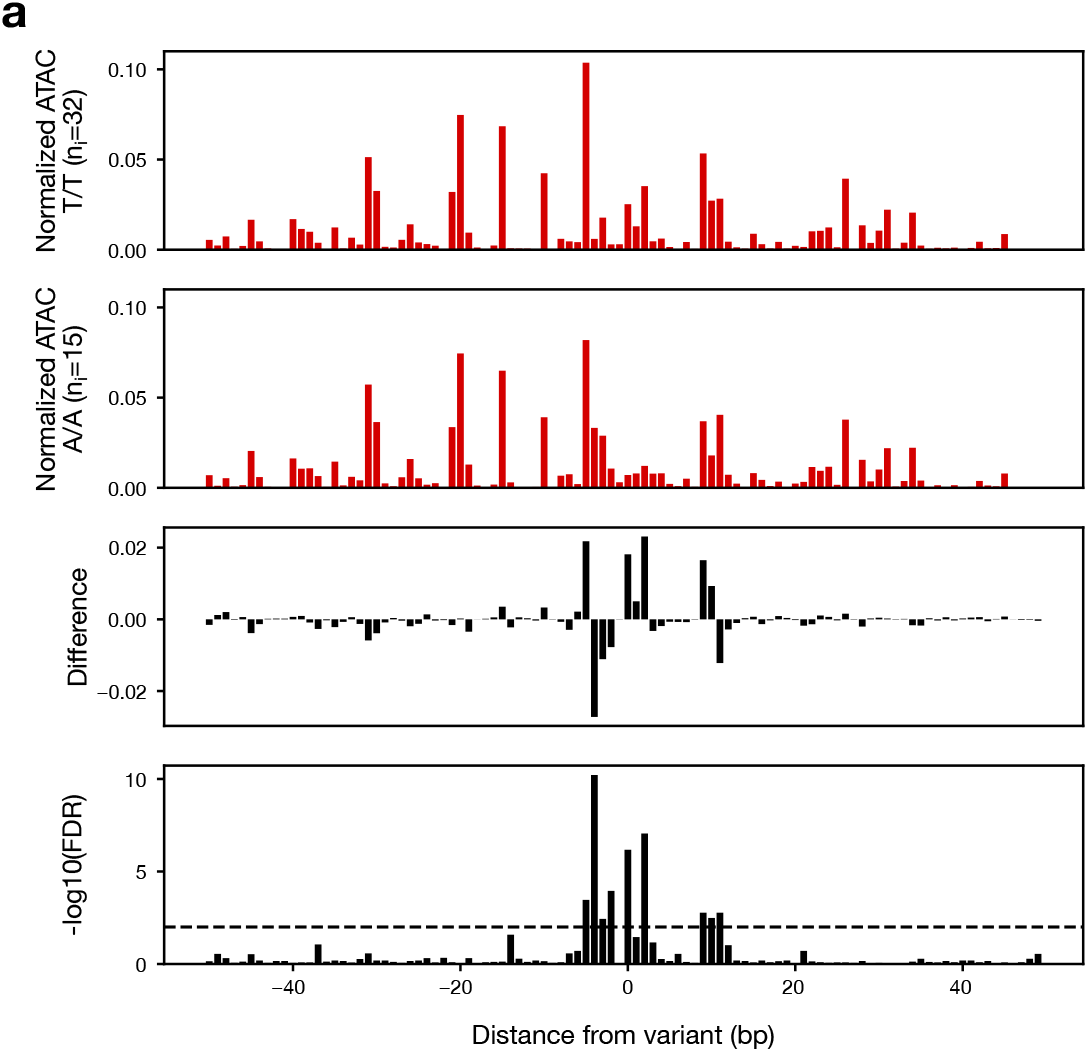
**a)** Extended version of Fig. 4a. Signal is shown for individuals from AFGR homozygous for the reference (T/T, 32 individuals) or alternate (A/A, 15 individuals) allele. The signal is normalized for read depth to emphasize shape differences. n_i_=number of individuals. The third panel shows the difference at each position between the two top panels. The last panel shows the Benjamini-Hochberg corrected FDR values for t-test comparisons done for each position. Dashed line denotes FDR=0.01.

## Methods

### Nona model

Nona is a multimodal masking based model that operates on DNA sequence and base-resolution tracks. At its core, Nona takes in masked DNA sequence and tracks, together with binary masks that specify which positions are masked, and predicts the masked values. In the following sections, we will explain the model in more detail.

#### Input and output

Nona operates on masked DNA sequence and base-resolution tracks. For a given model, the number of tracks T is fixed. The model takes as input a L_in_ × 4 masked one-hot DNA sequence and L_in_ × T matrix of masked tracks, where L_in_ is the input length. At masked positions in the sequence, all four channels are set to 0 (no information), while unmasked positions contain the standard one-hot base encoding. For each track, values are similarly set to 0 at masked positions and retain their observed values elsewhere.

In addition, Nona receives two binary mask inputs: one for sequence (L_in_ × 4) and one for tracks (L_in_ × T), each indicating which positions are observed (1) versus masked (0). For DNA sequence, the binary mask allows the model to distinguish between genuinely missing bases (“N”) and intentionally masked positions. All four sequence mask channels share the same value at a given position. Analogously, the tracks mask helps the model distinguish between true zeros and masked zeros. Thus, the cumulative input to Nona is a L_in_ × (2*(4+T)) matrix. In general, arbitrary bases in the sequence and pixels in the track matrix can be masked, allowing flexible masking schemes.

Given these inputs, Nona outputs an L_out_ × 4 matrix of predicted base probability logits, and an L_out_ × T matrix of predicted track signals on log scale, where L_out_ is the output length and L_out_ <= L_in_.

#### Architecture

The Nona architecture is inspired by the Borzoi model^2^ but adopts a simplified design with a constant channel width throughout. It follows a hybrid convolution–transformer–upsampling convolution U-Net structure. Sequence inputs are concatenated with their corresponding sequence masks, and track inputs are concatenated with their track masks. Separate convolutional encoders process the sequence and track inputs. The core reusable unit is a residual convolutional block consisting of a convolution, dropout, and batch normalization step with a skip connection.

The encoder path comprises an initial convolution followed by multiple reduction stages (n_reductions), each containing n_conv_per_reduction residual convolutional blocks and a max-pooling operation (stride 2) to downsample the representation. Before each pooling step, outputs are passed through a pointwise convolution and stored as skip connections for the decoder. The sequence and track encoder outputs are summed just before the transformer block. The transformer blocks are implemented using the x-transformers library, with flash attention and rotary positional embeddings enabled.

In the decoder, the representation is upsampled in n_reductions stages to recover the original sequence resolution. At each stage, the upsampled features are combined with the corresponding skip connections from both the sequence and track branches (if present) and passed through n_conv_per_reduction residual convolutional blocks per stage. Finally, the output passes through two additional convolutional layers, and the central L_out positions are cropped to match the desired output length. The final convolution output has 4 + T channels, where T is the number of tracks. This is split into two outputs corresponding to sequence and tracks. All output values are clipped to a fixed range (typically between -10 and ln(5000)) to stabilize training, and dropout is applied throughout the network to prevent overfitting.

During sequence-to-function training, where track inputs remain constant, the model explicitly zeros out the track skip connections and the deepest track representation to avoid batch normalization instabilities during inference. For functional language modeling and functional genotyping models, we use n_reductions=0, as that performed better than the U-Net versions.

#### Data loading and training loop

The data loader takes as input a reference genome, a set of genomic coordinates, a collection of unstranded base-resolution tracks, and user-specified mask definitions. To generate a single training example, a genomic locus is sampled from the provided coordinates. The coordinates are shifted by a “jitter” value drawn from Uniform(-J,J), where J is a user-defined maximum shift. The corresponding DNA sequence (L_in_ × 4) and track matrix (L_in_ × T) for the locus are then retrieved. During training, with 50% probability, the sequence is reverse-complemented and the tracks are reversed.

Given the mask specifications, a set of 4 masks is sampled: an input sequence mask (L_in_ ×, input track mask (L_in_ × T), output sequence mask (L_in_ × 4), and output track mask (L_in_ × T). For the input masks, 1 denotes observed and 0 denotes masked positions. For the output masks, 1 marks positions where the loss is to be computed. Importantly, the masks are sampled independent of the data. Details of the masking strategies and implementation are described in the next section.

Before model input, track values are transformed using log1p, and the sequence and tracks are multiplied by their respective input masks. The masked sequence, masked tracks, and corresponding input masks are provided to the model. The model outputs a L_out_ × 4 matrix of base probability logits, and an L_out_ × T matrix of predicted track signals. These predictions, together with labels cropped to L_out_ and the output masks, are retrieved for a mini-batch of inputs and the loss is computed. Both predicted and target tracks are clipped to a user-specified range before loss computation. The specific loss functions are described in the following section.

We train using the AdamW optimizer with weight decay 0. Learning rate schedules and other hyperparameters are specified for each model in their respective sections. We train with mixed precision (bfloat16) and use a gradient_clip_val of 0.5.

#### Masking implementation

The masking module defines how the DNA sequence and tracks are masked in the input and where the loss is computed in the output. Each sample from this module returns four binary masks: an input sequence mask (L_in_ × 4), input track mask (L_in_ × T), output sequence mask (L_in_ × 4), and output track mask (L_in_ × T). The module is initialized with a masking function, along with its associated parameters.

For the input masks, 1 denotes observed and 0 denotes masked positions. For the output masks, 1 marks positions where loss is computed. For the input and output sequence masks, all four mask channels share the same value at a given position. In most cases, the output masks are the logical inverse of the input masks; however, this is not enforced. For instance, in cases where certain parts of the data should be ignored entirely, both the input and output masks can be set to 0.

Masking schemes can be static or dynamic. In static masking, such as for sequence-to-function prediction, the DNA sequence is always fully observed while all tracks are masked. In dynamic masking, such as in masked language modeling, different subsets of positions are masked independently for each example. The fraction of positions to be masked and other options are specified as parameters to the masking function. The same genomic example can appear multiple times in training with different mask instantiations, improving generalization. During validation and testing, however, a fixed mask is used for each example to ensure reproducibility.

The framework provides several built-in masking functions and also allows custom user-defined masking strategies. No explicit constraints are placed on user-defined masking strategies beyond adhering to proper dimensions.

#### Loss

The model outputs an L_out_ × 4 matrix of predicted base probability logits, and an L_out_ × T matrix of predicted track signals. The signal tracks predictions are on log scale. During training, the data loader also provides corresponding binary output masks of the same dimensions to specify which positions contribute to the loss.

Two loss components are computed separately: one for sequence prediction and one for track prediction. The components are combined with a scalar weight applied to the sequence loss.

For each example, we compute a cross-entropy loss on the sequence, averaging over positions that are set to 1 in the output sequence mask, similar to discrete diffusion models for language^30^. The loss is multiplied by a scalar factor (seq_weight). For tracks, we use a Poisson negative log-likelihood loss, averaging over positions that are set to 1 in the output track mask. Each track can optionally be weighted by a corresponding coefficient (track_weights) to adjust for differences in signal scale or track importance.

Implementation details for both the sequence masked cross-entropy and Poisson losses are provided in the code below.

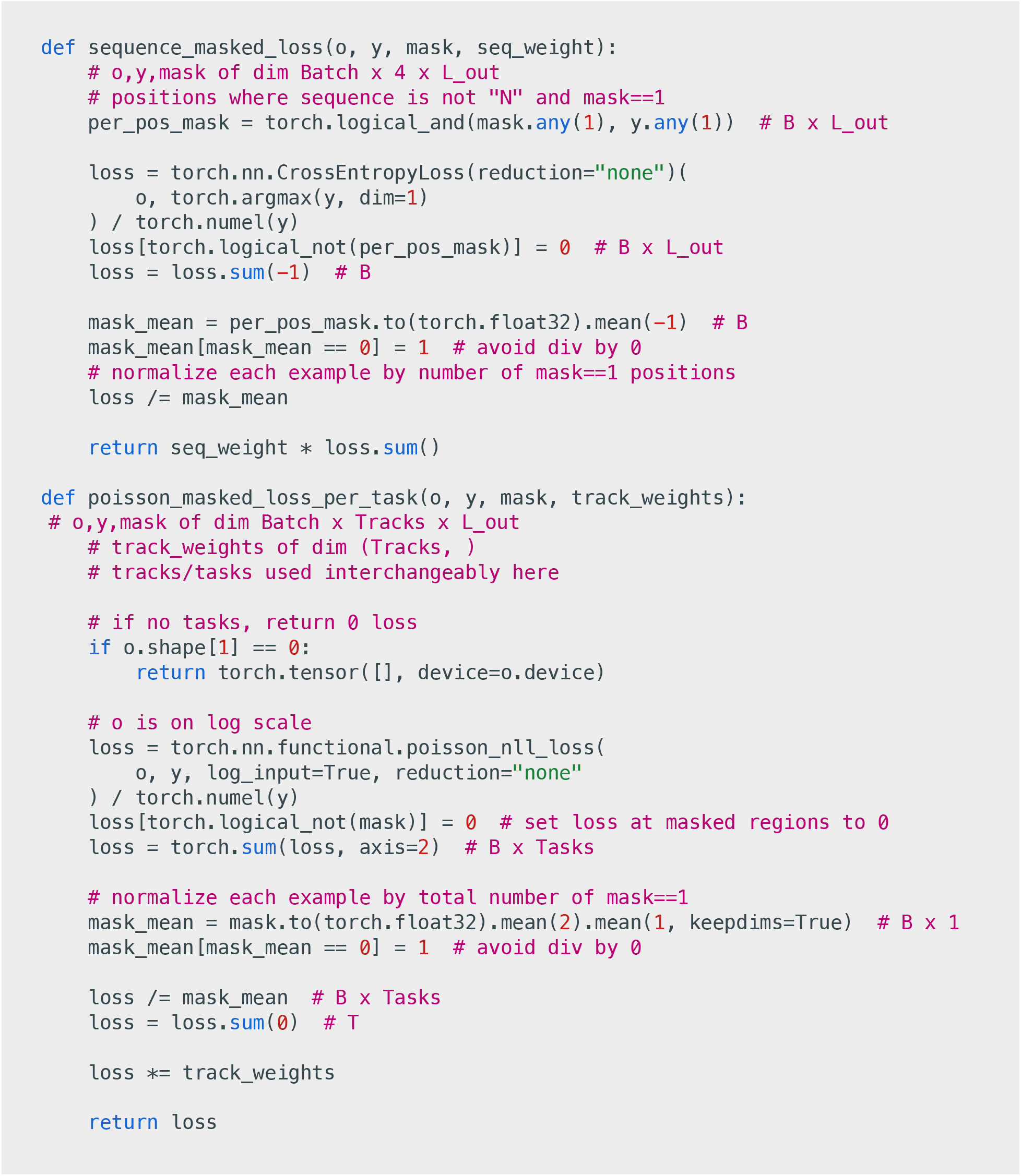

### Context-aware models

#### Model training

##### Data

We trained on both human and mouse data, using a subset of Enformer’s tracks^20^. For human, we included 7 K562 DNase-seq tracks and 2 K562 CAGE-seq tracks. For mouse, we used 4 DNase-seq (3 from MEL cells and 1 from leukemia stem cells) and 3 CAGE-seq tracks from hematopoietic or myeloid progenitor cell types, selected as the closest available analogs to K562.

For DNase-seq tracks, we downloaded the read-depth normalized signal track BigWigs from the ENCODE portal and merged replicates, if any. For CAGE-seq, we downloaded the forward and reverse strand BigWigs from the FANTOM portal and merged them to make unstranded tracks. Replicates were not considered. Track identifiers are provided below.

Human, DNase-seq: ENCFF899YDP. ENCFF515UNC, ENCFF708UIS, ENCFF413AHU, ENCFF868NHV, ENCFF565YDB, ENCFF971AHO

Human, CAGE-seq: CNhs11250, CNhs12336

Mouse, DNase-seq: ENCFF990ATO, ENCFF223QRV, ENCFF155MQS, ENCFF014NWA

Mouse, CAGE-seq: CNhs12202, CNhs12198, CNhs12551

We used the same train, validation and test splits as Enformer for both human and mouse, with 196608 bp input length and 114688 bp output length. We used a jitter value of 3 bp for the initial sequence-to-function pre-training. For the context-aware fine-tuning, we used a jitter of (114688-4096)/2 = 55296 bp. This ensures that the labels come from the same regions as the pre-training. Batches consist of a mix of human and mouse examples.

##### Architecture

The architecture uses 512 channels, n_reductions=7, n_conv_per_reduction=2, with kernel size 7, and 12 transformer layers with 8 attention heads each. The model has input length of 196608 bp. For the sequence-only model, the output is cropped to 114688 bp, while for the context-aware model, the output is cropped to 4096 bp. The architecture has ∼120M parameters. The outputs are clipped between (-10, 8.517).

##### Masking

For the sequence-to-function pre-training, we used a static input mask in which the 196608 bp sequence is fully presented while all tracks are fully masked. For the output mask, for human examples, the loss was computed over the human tracks, and similarly for mouse examples.

For the context-aware fine-tuning, we use a mask in which the sequence is fully presented. For human examples, the mouse tracks are fully masked (as they are unavailable), and human tracks are masked in the central 4096 bp but presented everywhere else (i.e. 196608-4096 = 192512bp), and similarly for mouse examples. For the output mask, for human examples, the loss is computed in the central 4096 bp over human tracks, and similarly for mouse examples.

##### Training details

For the initial sequence-only training, we trained 5 model replicates with the same training splits, but 5 different random seeds. Each model was trained for 150,000 steps with a fixed learning rate of 1×10^-4^ and batch size of 64. Training each model took 1.5 days on 8 NVIDIA B200 GPUs. Models with the best log1p transformed Pearson correlation at 128 bp resolution on human tracks on the validation set were saved.

For the context-aware fine-tuning, we initialized 5 models using the best sequence-only models from each seed. The optimizers were reinitialized, and models trained with a fixed learning rate 1×10^-4^. Each model was trained for 100,000 steps with a batch size of 128. Each model took 1 day to train on 16 NVIDIA B200 GPUs. Models with the best log1p transformed Pearson correlation at 128 bp resolution on human tracks on the validation set were saved.

The models were trained using PyTorch v2.7 and PyTorch Lightning v2.5.3.

#### Test set evaluation

We evaluated both the sequence-to-function and the context-aware models on the test set. While both models take a 196608 bp window as input, the sequence-to-function model outputs predictions for the central 114688 bp while the context-aware model outputs predictions for the central 4096 bp. To ensure comparable evaluation, we divided each 114688 bp test region into 4096 bp non-overlapping chunks (yielding 28 chunks per region). For each chunk, we ran both models centered on the chunk. We extracted the central 4096 bp prediction for the sequence-to-function model, and the corresponding 4096 bp prediction for the context-aware model. In this way, we obtained predictions over the entire test set. Both predictions and labels were aggregated at 128 bp resolution and log1p transformed before computing downstream metrics.

For comparison against Enformer predictions, we accessed the model through gReLU^39^. We used the matching track outputs from Enformer after adjusting for track-specific normalization factors.

#### ChromHMM analysis

To identify loci where the context-aware model made distinct predictions, we compared its outputs to those of the Nona sequence-only model and Enformer. We defined “outlier” regions with increased differential predictions (context-up) as 4096 bp windows where the log2 fold change (log2FC) between Nona context-aware and Nona sequence-only predictions exceeded 1, and the log2FC between Nona context-aware and Enformer predictions also exceeded 1. Conversely, regions with decreased differential predictions (context-down) were defined as those with log2FC < –1 in both comparisons.

All analyses were restricted to test-set regions. As a background, we used all test-set regions. chrX regions were excluded from the analysis.

Chromatin-state annotations were obtained from the EpiMap project^40^ (track BSS01057, “CA.ML.K562,” 18-state ChromHMM model; https://personal.broadinstitute.org/cboix/epimap/ChromHMM/observed_aux_18_hg38/CALLS/BSS01057_18_CALLS_segments.bed.gz). For each set of regions (context-up, context-down, and background), we computed the fraction of bases overlapping each ChromHMM state.

#### ChromHMM visualization at example locus

To visualize ChromHMM states at the example locus, we used the Epilogos browser^23^. We used the Boix *et al*., 18-state model and restricted it to Immune samples (URL: https://epilogos.altius.org/?application=viewer&sampleSet=vC&mode=single&genome=hg38&model=18&complexity=KL&group=Immune&chrLeft=chr7&chrRight=chr7&start=57393464&stop=57508152&gatt=cv).

#### TRIP-seq analysis

##### Datasets

###### Leemans et al. 2019

We obtained TRIP-seq data from Table S1 and Dataset S2 of Leemans *et al*. 2019^21^. We extracted the seven promoter sequences from hg38, accounting for their annotated strand orientation. Each example was reconstructed by concatenating the promoter sequence, the GFP reporter sequence (https://www.ncbi.nlm.nih.gov/nuccore/KC710228.1), the unique barcode, and the sNRP-1 polyA sequence (AAATAAAATACGAAATGA). The resulting construct was inserted into the corresponding genomic integration position in a strand-aware manner.

###### Hong et al. 2022

We obtained the data from https://zenodo.org/records/7076228 as processed by Karollus *et al*.^41^, and adapted the preprocessing code from https://github.com/Karollus/ SequenceModelBenchmark. The data was originally generated by Hong *et al*.^22^. For each example, we concatenated the promoter sequence with the tdTomato gene sequence (in “minimal” configuration) and inserted the construct into the integration position, again in a strand-aware manner.

#### Predictions and evaluation

All evaluations used predictions from the CNhs12336 CAGE-seq track (following Karollus *et al*.^41^), corresponding to K562 CAGE-seq signal. Predictions from the Nona models were aggregated to 128 bp resolution to match Enformer’s output binning. For each construct, the predicted signal in the bin containing the transcription start site (TSS) and the two neighboring bins (± 1 bin) were summed to yield a single promoter-level prediction.

Following Karollus *et al*.^41^, predictions were averaged across both forward and reverse sequence orientations and over three TSS jitter offsets (−43 bp, 0 bp, +43 bp), resulting in six configurations per construct. The 0 bp offset corresponds to placing the TSS at the center of the input window. Model replicates were evaluated individually and also as ensembles formed by averaging replicate predictions.

For context-aware predictions, the observed tracks were extracted over bases corresponding to genomic coordinates. Any values in the central 4096 bp (after inserting the reporter sequence) were masked.

Performance was quantified using Spearman correlation between observed and predicted expression levels across genomic integration sites separately for each promoter.

### Functional language modeling

#### Model training

##### Data

We downloaded the GM12878 DNase-seq read-depth normalized BigWig track (accession ENCFF093VXI) from the ENCODE portal. We downloaded the peak file (accession ENCFF588OCA) corresponding to the same experiment. We created a set of GC-matched, no peak regions using the get_gc_matched_intervals function in the gReLU package^39^. We split our data into the following splits based on chromosomes: chr1: test, chr8,chr10: validation, rest in training. Training was performed on the union of peak and non-peak sequences. We used a jitter value of 512 bp during training.

##### Architecture

We use n_reductions=0, thus the architecture did not consist of pooling/upsampling blocks and skip connections. We observed this performed better than U-Net based architectures for this application. The model has input and output length 1024 bp. Our model uses 512 channels, 20 transformer layers each with 8 attention heads. The architecture has ∼64.8M parameters. The outputs are clipped between (-10, 8.517).

##### Masking

For functional language modeling, we use a mask in which the track is always provided unmasked as an input to the model. For the sequence, we first sample a masking rate r from the Uniform(0,1) distribution, and use that as the rate for masking each base independently^30^. In case no base is masked, we resample r and repeat the procedure till at least one base is masked.

##### Training details

We train two separate Nona models. One of the models consists of one track, while the other uses no tracks (and thus is a pure masked language model). The rest of the details are identical for both models.

We trained each of the models for 1,000,000 steps with a batch size of 512. We used a learning rate schedule consisting of a linear warmup for the first 2000 steps up to a learning rate of 3×10^-4^ followed by cosine decay over the next 998,000 steps. Training took 2.5 days for each model on 8 NVIDIA B200 GPUs.

The models were trained using PyTorch v2.7 and PyTorch Lightning v2.5.3.

#### Importance scores

##### Nona

To extract importance scores from a conditional masked language Nona model, we first tried an approach where we mask one base at a time and look at the predicted distribution over A,C,G, and T for each base. Thus, for a sequence of length L, we make L predictions through the model to obtain an importance score for every base in the sequence. We convert the predicted distribution to an importance score by computing the negative of the entropy (base 2). Higher scores indicate greater model confidence at that base. Prior to visualization, we subtract the median per-base importance score computed across a set of sequences.

As this procedure is slow, we also examined a single-pass approach, where we provide the model with all tracks and all bases simultaneously– the input is fully unmasked. Strictly speaking, the model was not trained on such inputs, as it was never trained to reconstruct any base that was unmasked in the input. Empirically, we observed that the per-base importance scores obtained in a single-pass correlated strongly with the one-by-one inference method outlined above. Thus, for the results in the paper, we derived importance scores using the single-pass approach.

##### Caduceus

We accessed the trained Caduceus^26^ model from HuggingFace (model: kuleshov-group/caduceus-ph_seqlen-131k_d_model-256_n_layer-16). We loaded the model using the AutoModelForMaskedLM class from HuggingFace’s transformers library v4.55.0.

Unlike Nona, Caduceus is trained with a fixed masking rate based on the masking recipe in BERT^42^. Nonetheless, we observed that masking one base at a time resulted in reasonable importance scores. However, the single-pass approach outlined above correlated poorly with the one-by-one approach. To address this, we developed a balanced masking batch procedure to obtain more stable per-base predictions.

In this approach, each input sequence is replicated V times to form a batch. In each replicate, bases are randomly masked at rate p, but the masking is arranged such that across the entire batch, every position is masked exactly the same number of times (V*p). This guarantees balanced coverage of all positions while preserving the model’s training regime. Predicted probabilities are aggregated across the V*p replicates for each position. Finally, we convert these distributions into importance scores as described in the Nona section above. This balanced masking batch method provides deterministic, reproducible per-base scores that correlate well with the one-by-one masking approach but are much faster to compute. We used V=20,p=0.15.

##### Sei

To obtain importance scores for a sequence-to-function model, we downloaded the Sei model^28^ and weights from https://zenodo.org/records/7943307. We loaded the model as sei = NonStrandSpecific(Sei(4096, 21907)), but only used sei.model for importance score calculation. We used the deep_lift_shap function in tangermeme (v0.5.1)^43^ using the average predictions across all of Sei’s GM12878 DNase-seq output tracks as the target. Since our sequences are of length 1024 bp while Sei expects a 4096 bp input, we embedded our sequences in a 4096 bp sequence centered at hg38 chr4:19599500, which is inaccessible in GM12878, and cropped the final scores back to 1024 bp around our sequence of interest.

##### Enrichment analysis within repeats and motifs

After computing importance scores for the different models, our goal was to analyze enrichment of the masked language models’ importance scores within repeat elements and transcription factor motifs. To get repeat element annotations, we downloaded the https://www.repeatmasker.org/genomes/hg38/RepeatMasker-rm405-db20140131/hg38.fa.out.gzfile from the RepeatMasker^29^ website on 2025-09-16. We considered any base that fell within any element annotated in this list to be within a repeat element. We computed importance scores for Nona trained with GM12878, Nona trained without any track, Caduceus, and Sei within GM12878 DNase-seq peaks in the test set of the Nona models. We considered bases within the top 1% of Sei importance scores and not within a repeat element as a proxy for putative transcription factor binding sites.

As the importance scores from different masked language models are not directly calibrated, we rank-normalized the values: each model’s importance score was replaced by its empirical rank (computed across all sequences and positions) and rescaled to the [0,1] interval, ensuring comparability across models. Finally, we looked at the distribution of each of the model’s importance scores within the bases marked in the repeat and motif sets.

### Conditional sequence design

#### Generation algorithm

We use the functional language model to design sequences that satisfy specified track profiles. The input (“prompt”) consists of the desired profiles across all tracks, a sampling temperature parameter T, and a parameter B specifying how many bases to fill per step. The track profiles are always provided as input to the model. Sequence generation proceeds in multiple rounds, with a fixed number of new bases filled each time until the sequence is complete. At the start, the sequence is masked at all positions.

In each round, the model produces logits (pre-softmax scores) for every masked position. These logits are divided by the temperature T before applying the softmax, which controls the sharpness of the probability distribution. Lower values of T concentrate probability mass on the most likely base, while higher values distribute it more evenly across alternatives. In general, lower temperatures result in higher quality but biased samples. We use T=0.7 by default.

From the currently masked positions, a subset of size B is chosen at random. For each selected position, a base is sampled independently according to the temperature-scaled probability distribution. Once filled, these positions remain fixed for the rest of the process.

#### Oracle-based evaluation of generated sequences

We generated sequences by conditioning on the profiles of held-out test sequences. To evaluate the generated sequences, we compared the predictions of an oracle model on the original test sequence to the matched generated sequence, following previous work^11,12^.

##### Oracle

We used fold0 of the Borzoi model^2^ as an oracle, accessing it through gReLU^39^. Borzoi returns predicted profiles at 32 bp resolution. Since Borzoi is trained on long sequences (512kb), and our designed sequences are short (1024 bp), we embed the short sequences in longer background sequences before running Borzoi on them and crop the output corresponding to the 1024 bp sequence. We observed that averaging over 5 background sequences of length 32kb provided a sufficient tradeoff between accuracy and inference time. We used the prediction corresponding to the ENCFF093VXI track in Borzoi (GM12878 DNase-seq, same track as the one used by the Nona model).

##### Metrics

For each genomic sequence in our evaluation set, we generate a sequence using Nona (or a baseline method) by conditioning on its profiles. We then run the oracle on the genomic sequence and the corresponding generated sequence and compare the predictions. The oracle outputs a predicted profile vector of length P for each sequence. Thus for N sequences, we obtain two NxP matrices from the oracle: one for the genomic (R) and one for the matched generated sequences (G). We use the following metrics:

1. Mean-squared error (MSE): mean((R-G)^2^) directly measures agreement between predicted profiles of genomic and generated sequences. Lower is better.
2. Total counts correlation: spearman(R.sum(-1),G.sum(-1)) measures the correlation between total predicted counts across sequences. Higher values indicate better quantitative control of overall signal strength.
3. Total shape correlation: mean([spearman(R[i], G[i]) for i range(len(R))]) measures the similarity of predicted profile shapes on a per-sequence basis, independent of scale. Higher is better.
4. Predictive distribution KS distance: KS(R.sum(-1), G.sum(-1)) computes the Kolmogorov–Smirnov statistic between the distributions of total predicted counts for genomic versus generated sequences. Lower values indicate the generated sequences better match the distributional properties of the genomic sequences.
5. K-mer spectrum shift: Following Sarkar *et al*.^11^, we compare the distribution of short sequence motifs between genomic and generated sequences, independent of oracle predictions. Specifically, we compute normalized frequency spectra of all possible 6-mers and measure the divergence between genomic and generated spectra using the Jensen–Shannon distance. Lower values indicate that the generated sequences more closely match the k-mer composition of the genomic sequences.

##### DDSM

We obtained the source code for DDSM^12^ from its official GitHub repository (https://github.com/jzhoulab/ddsm/) and followed the described training setup. The model employs a Jacobi diffusion process with parameters s=2/(a+b), where a=1 and b=3 as recommended by the authors (optimal b is k-1, where k is the number of categories) and a maximum diffusion time of 4.

We used the custom-designed 1D convolutional architecture introduced in the original DDSM paper. The model comprises 20 layers of 1D convolutions, interleaved with time-embedding and normalization layers. Training was performed for 200 epochs with a learning rate of 5×10^-4^ using early stopping based on the mean squared error (MSE), evaluated using Borzoi’s ENCFF093VXI task as the oracle. We used a batch size of 128, and Adam with default parameters. Training took around 57 hours on 1 NVIDIA A100 GPU.

For sampling, we used the Euler–Maruyama sampler to generate sequences from the trained model. Generated samples were discretized using an argmax operation over k categorical dimensions; this mapping is effectively deterministic, as trained model outputs are typically near 1 for one category and near 0 for others.

##### Evaluation

We consider a random subset of 10,000 test sequences (both peaks and non-peaks) for evaluation. For each sequence, we provided its corresponding profile as input to a design method, and generated one sequence per method. Metrics were computed using Borzoi as an oracle as described above for each pair of genomic and generated sequences. For Nona with GM12878, we varied the temperature and bases per step parameters. For DDSM, we varied the number of steps and time dilation parameters.

### Functional genotyping

#### Data processing

##### Tracks and training regions

From ENCODE accession ENCSR637XSC (GM12878 ATAC-seq), we downloaded 3 BAM files: ENCFF440GRZ, ENCFF962FMH, ENCFF981FXV. We merged the BAM files in one. To convert to a base-resolution BigWig file, we used the reads_to_bigwig.py script from ChromBPNet^3^. The analysis was performed within a Docker container instantiated from the kundajelab/chrombpnet:latest image (digest sha256:c2f470debe75113652311b131eed2894ec5354694a071157c0825c49366b5f29). This BigWig file was used for training. We used the same training regions and splits as for the GM12878 DNase-seq functional language model. We used a jitter value of 500 bp for augmentation.

##### AFGR ATAC-seq and genotypes

###### ATAC-seq

We downloaded ATAC-seq BAM files corresponding to 100 individuals from the African Functional Genomics Resource (AFGR) project^33^ from the ENCODE portal in January 2025, with API request https://www.encodeproject.org/search/? searchTerm=AFGR&type=Experiment&limit=200. For subsampling experiments, we subsampled each BAM to depths of 1M, 2M, 3M, 4M, 5M, and 10M using samtools view -s. We converted each of the original and subsampled files to separate BigWig files using the reads_to_bigwig.py script from ChromBPNet. The created BigWig files contain base-resolution signals where the value at a specific base corresponds to the number of reads in the BAM file that either start or end at that position, after accounting for Tn5 shift. Thus, they do not explicitly contain any read level information.

###### Genotypes

We first identified a set of common SNPs within regions with high ATAC-seq signal in LCLs. To do this, we first obtained common SNPs from the UCSC Genome Browser *snp151Common* track. The full table was downloaded from: https://hgdownload.soe.ucsc.edu/goldenPath/hg38/database/snp151Common.txt.gz. We restricted ourselves to single-nucleotide variants. We overlapped these SNPs with the top 1,000 ATAC-seq peaks ranked by signal strength from an LCL sample (GM18498; peak file ENCODE accession ENCFF717TCQ).

Genotypes for these SNPs were retrieved from the 1000 Genomes Project high-coverage dataset (accession: 1000G_2504_high_coverage, 20201028 release). For each SNP locus, we extracted genotype information from the per-chromosome VCFs (ftp://ftp.1000genomes.ebi.ac.uk/vol1/ftp/data_collections/1000G_2504_high_coverage/working/20201028_3202_raw_GT_with_annot/), using tabix to fetch the region and bcftools to filter single-base alleles.

In total, we obtained genotypes at 4756 SNPs across 3202 individuals, of which 83 overlapped with the AFGR ATAC-seq dataset. All processing was performed in January 2025.

#### Model training

##### Architecture

We use n_reductions=0, thus the architecture did not consist of pooling/upsampling blocks and skip connections. The model has input and output length 128 bp. Our model uses 256 channels, 8 transformer layers each with 8 attention heads. The architecture has ∼8.8M parameters. The outputs are clipped between (-10, 8.517).

##### Masking

We used the same mask as for the functional language model.

##### Training details

We trained a single model on the GM12878 base-resolution ATAC-seq. We trained the model for 250,000 steps with a batch size of 256. We used a fixed learning rate of 3×10^-4^. Training took 6 hours on 1 NVIDIA A100 GPU. The models were trained using PyTorch v2.2 and PyTorch Lightning v2.5.0.post0.

#### Evaluation

##### GM12878 evaluation

We evaluated whether the model could correctly predict GM12878’s alleles at common SNPs. Evaluation was performed on the 4756 SNPs in the top 1000 ATAC-seq peaks described in the Genotypes section. To avoid ambiguity, we restricted to sites where GM12878 is homozygous for either the reference or alternate allele. We further stratified SNPs by whether they overlapped the training or held-out test regions. Because the model was trained on the hg38 reference genome rather than GM12878’s genome, sites where GM12878 is homozygous alternate represent cases where the model is systematically trained toward the wrong base.

To extract the model’s prediction, we input the base-resolution ATAC-seq track centered at the variant, along with the adjacent sequence, masking the focal base. A prediction was counted as correct if the base with the highest predicted probability among A,C,G,T matched GM12878’s true allele.

##### Log-likelihood score for linking

We next considered whether an individual’s base-resolution ATAC-seq BigWig could be linked back to their genotype in a linking (re-identification) attack^34^. The inputs to this analysis are a BigWig file from one individual and a panel of genotypes from multiple individuals, one of whom is the true source of the BigWig. Our goal is to determine whether the BigWig can be correctly linked to its source individual.

To score the match, we used a simple log-likelihood approach. Using the model, we obtained predicted base probabilities for a set of SNPs. For a given individual, we computed a per-SNP log-likelihood based on their genotype:

- homozygous reference: 2*log(p(ref))
- homozygous alternate: 2*log(p(alt))
- heterozygous alternate: log(p(ref))+log(p(alt))

Here p(ref) and p(alt) are the model’s predicted probabilities at the reference and alternate bases. We renormalize them by omitting the other 2 bases, such that p(ref)+p(alt)=1 for each SNP. The total log-likelihood score for each individual is the sum across SNPs to obtain a linking score, and the individual with the highest score is predicted as the source of the BigWig.

##### GM12878 linking analysis

We applied the log-likelihood method to the GM12878 base-resolution ATAC-seq data. Model predictions were obtained at the 4756 SNPs in highly accessible regions, and linking scores were computed for each of the 3202 individuals in the 1000G dataset.

##### AFGR linking analysis

We extended the analysis to 83 ATAC-seq samples from the AFGR project. Strictly speaking, a separate model should be trained for each sample because of differences in sample quality and sequencing depth. However, we observed that reusing the model trained on GM12878 without any read depth normalization still achieved high linking performance. This indicates that the model relies primarily on base-resolution profile shape rather than read depth for genotype inference.

For each AFGR sample, we computed linking scores against all 3202 individuals in the 1000G dataset. A linkage was marked correct if the true source individual of the ATAC-seq file had the highest score. We repeated the analysis on subsampled ATAC-seq data at depths of 1M, 2M, 3M, 4M, 5M, and 10M reads.

